# OpenSim Moco: Musculoskeletal optimal control

**DOI:** 10.1101/839381

**Authors:** Christopher L. Dembia, Nicholas A. Bianco, Antoine Falisse, Jennifer L. Hicks, Scott L. Delp

**Affiliations:** Department of Mechanical Engineering, Stanford University, Stanford, California, United States of America; Department of Movement Sciences, KU Leuven, Leuven, Belgium; Department of Bioengineering, Stanford University, Stanford, California, United States of America; Department of Orthopaedic Surgery, Stanford University, Stanford, California, United States of America

**Author notes:** These authors contributed equally to this work (Christopher L. Dembia, Nicholas A. Bianco).

## Abstract

Musculoskeletal simulations of movement can provide insights needed to help humans regain mobility after injuries and design robots that interact with humans. Here, we introduce Open-Sim Moco, a software toolkit for optimizing the motion and control of musculoskeletal models built in the OpenSim modeling and simulation package. OpenSim Moco uses the direct collocation method, which is often faster and can handle more diverse problems than other methods for musculoskeletal simulation but requires extensive technical expertise to implement. Moco frees researchers from implementing direct collocation themselves, allowing them to focus on their scientific questions. The software can handle the wide range of problems that interest biomechanists, including motion tracking, motion prediction, parameter optimization, model fitting, electromyography-driven simulation, and device design. Moco is the first musculoskeletal direct collocation tool to handle kinematic constraints, which are common in musculoskeletal models. To show Moco’s abilities, we first solve for muscle activity that produces an observed walking motion while minimizing muscle excitations and knee joint loading. Then, we predict a squat-to-stand motion and optimize the stiffness of a passive assistive knee device. We designed Moco to be easy to use, customizable, and extensible, thereby accelerating the use of simulations to understand human and animal movement.

## Introduction

Musculoskeletal simulations have shed light on movement disorders by, for example, discovering ways to walk that reduce knee loading [1], revealing that children with cerebral palsy exhibit simplified motor control when walking [2], and reproducing eye disorders that cause double vision [3]. Simulations provide insights to increase safety as we push the limits of human performance, whether that is designing exercise equipment for preserving bone density in low gravity [4], designing exoskeletons to reduce back injuries from heavy lifting [5], or assessing the effect of training programs on knee ligament injuries in soccer [6]. Beyond human health and performance, researchers use musculoskeletal simulations to understand how animals move, for example by studying dinosaur locomotion [7] and differences between human and chimpanzee strength [8].

Simulations of movement are often categorized by whether the motion is prescribed from data or predicted by the simulation. One may prescribe the motion of a musculoskeletal model [9, 10] to estimate unmeasured quantities such as muscle-level energy consumption [11,12]. Motion prediction [13] can establish cause-effect relationships and help design clinical interventions, such as discovering gait impairments that arise from weakness and contracture [14] and designing prostheses for amputees [15]. Existing prescribed motion methods are rapid but either not customizable (e.g., cost functions are predefined) or lack support for important model features such as muscle dynamics. Predicting a motion often requires using a model with simplified musculature, reducing the dimensionality of the control scheme, or waiting many hours or days for a solution. A third category, which lies between prescribing and predicting, is tracking a motion, where errors between experimental and simulated kinematics are minimized [16]. To accelerate the application of simulations to scientific and clinical questions, researchers need unified simulation tools for solving diverse problems that span the prescribed-to-predicted spectrum and involve a variety of costs and unknown model parameters. For example, clinicians may be interested in subtle changes to an observed motion that minimize joint loading [1], and device designers may seek optimal stiffness parameters for a passive exoskeleton to improve the speed of an observed motion, but off-the-shelf simulation tools cannot handle these problems.

Most musculoskeletal simulation problems are naturally posed as optimal control problems: we seek a system’s parameters and time-varying controls that minimize a cost (e.g., energy consumption) subject to the dynamics of the system, expressed as differential-algebraic equations. An increasingly popular method for solving optimal control problems is to approximate the system’s states and controls as polynomial splines and solve for the knot points that lead the spline to obey the system’s dynamics [17–20]. The dynamics are enforced by requiring the time derivative of the state splines to match the derivative from the system’s differential equations at specified time points. This method is called “direct collocation” because the spline derivatives are “collocated” with the exact derivatives ([21], page 211; [22], page 498). Direct collocation produces a nonlinear program in which the system states are introduced as variables and the system’s dynamics are enforced as constraints. Typical musculoskeletal models lead to optimization problems with thousands of variables, yet these problems are tractable because the constraints enforcing dynamics at a given time depend only on the variables near that time; this speeds optimization. The method can handle a wide range of objectives and optimize model parameters, and leads to nonlinear programs that can be solved by generic optimization software.

The advantages of direct collocation have led biomechanists to use the method for tracking motions [16,23], predicting motions [24–33], fitting muscle properties [34], and optimizing design parameters [35]. Researchers have made key methodological advances, including efficiently handling multibody and muscle dynamics via implicit formulations [36, 37], minimizing energy consumption [38, 39], and employing algorithmic differentiation to simulate complex models more rapidly compared to using finite differences [40]. Along with these methodological advances, researchers have discovered that minimizing an energy-related cost produces non-physiological motions during walking [24], skipping is the most efficient gait on our moon [41], and unilateral amputees can improve gait symmetry with only a minor increase in effort [42].

Despite its advantages, direct collocation is not easy to implement, and very few laboratories have been able to apply this powerful technique. The method requires arduous bookkeeping of variables and efficient calculation of the objective and constraint function derivatives required by gradient-based optimization algorithms. Several direct collocation solvers exist (e.g., [43, 44]), but current solvers require users to incorporate their musculoskeletal models manually. OpenSim is a software package used by thousands of biomechanics researchers to model musculoskeletal systems [45–47], but OpenSim does not provide direct collocation. Several biomechanists have graciously shared their code combining OpenSim or hand-coded models with direct collocation solvers, but such code is tailored to specific models or motions, handles only unconstrained models, contains closed-source components, or is difficult to install on one’s own computer. Lastly, choosing the problem formulation (e.g., expressing dynamics as explicit or implicit differential equations) and solver settings (e.g., detecting sparsity patterns automatically) that lead to fast convergence requires expertise; ideally, such expertise is embedded into the software via defaults, and users can edit their formulation or solver settings with single commands.

To improve the accessibility of advanced optimal control methods in musculoskeletal biomechanics, we introduce OpenSim Moco (“musculoskeletal optimal control”), an easy-to-use, customizable, and extensible software toolkit for solving optimal control problems with OpenSim musculoskeletal models. OpenSim frees biomechanists from implementing equations of motion on their own, and OpenSim Moco frees biomechanists from implementing direct collocation. Moco not only removes the need to set up gradient-based optimization, but also includes an interface that abstracts away the details of constructing an optimal control problem. With just a few lines of code, researchers can solve complex problems with an OpenSim model for almost any movement. Users can add custom cost terms or constraints if Moco does not provide what they need. This paper first details the design and implementation of the software. We next show that Moco can accurately estimate muscle activity for an observed walking motion and allows users to customize the cost terms that are minimized. Finally, to illustrate that Moco can rapidly predict motions and model parameters, we predict a squat-to-stand motion and optimize the stiffness of a passive assistive device.

## Design and implementation

Moco solves optimal control problems that users define using a library of cost and constraint modules, which are implemented through configurable software classes. Users describe their problem with the *MocoProblem* class. To decouple the problem from the numerical methods used to solve it, we use the *MocoSolver* class. (We denote names of classes in Moco and Moco’s dependencies with italics.) Moco classes are available via C++, MATLAB, Python, and XML text files, with interfaces familiar to OpenSim users. We package the *MocoProblem* and *MocoSolver* together into a *MocoStudy* (Figure 1), which can be written to and loaded from XML text files. Moco contains utilities for visualizing and plotting a study’s solution, which is held by the *MocoSolution* class. For certain standard biomechanics problems, Moco provides simpler interfaces that may be preferable to the flexibility of *MocoStudy*.

**Figure 1.**
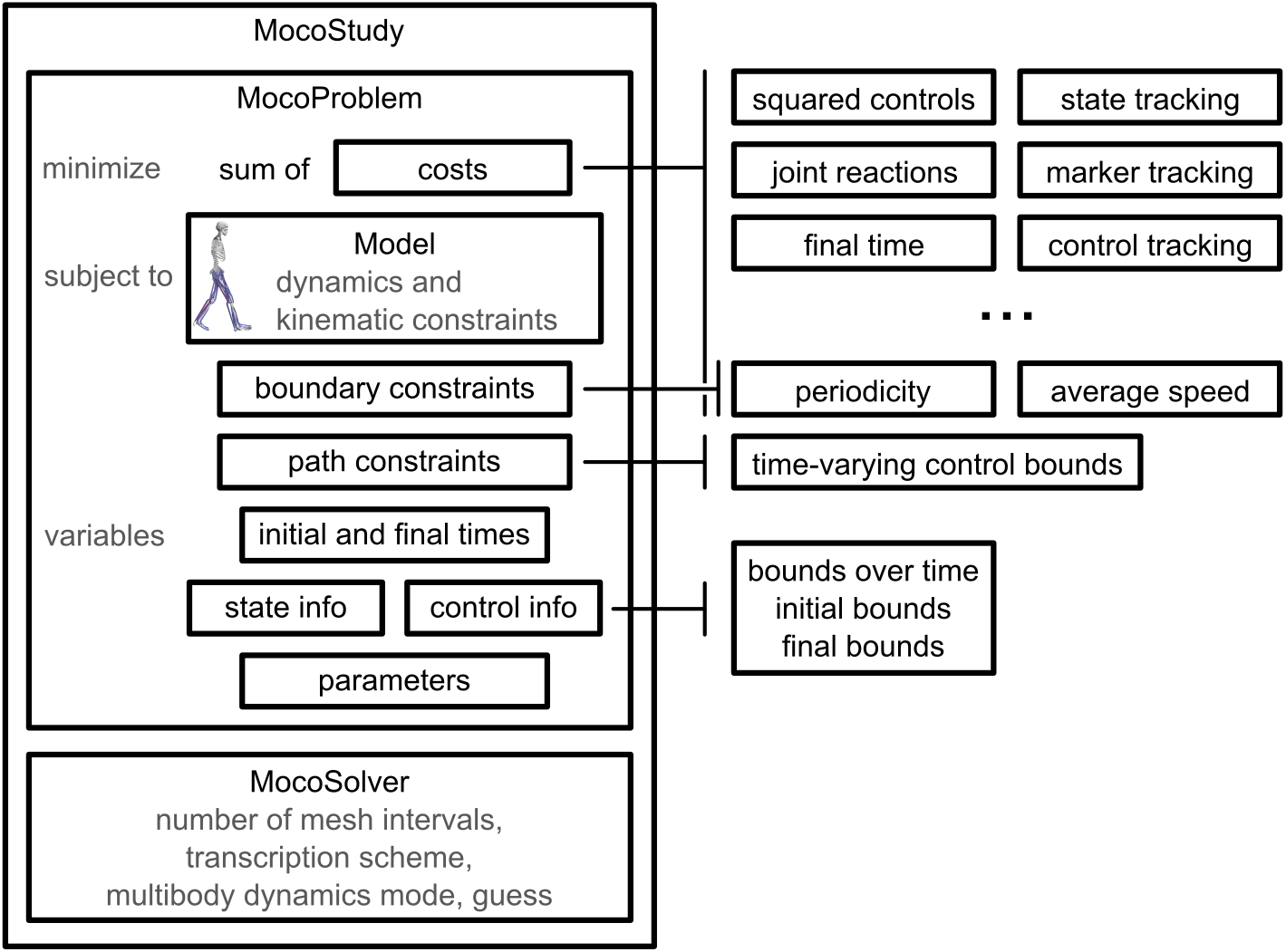
Overview of *MocoStudy*. Moco can solve custom optimal control problems using a library of cost, boundary constraint, and path constraint modules. Moco contains additional cost modules beyond what is shown here, and users can define their own custom modules.

### Defining problems with *MocoProblem*

*MocoProblem* supports a diversity of scientific questions and contains the following elements.

- **cost terms**: Users can minimize a weighted sum of control effort, deviation from an observed motion, joint reaction loads, the duration of a motion, and other costs by appending to the *MocoProblem* an instance of the class associated with a cost module (e.g., *MocoControlGoal* implements the control effort cost).
- **multibody dynamics, muscle dynamics, and kinematic constraints**: OpenSim *Models* are a standard way to describe musculoskeletal systems, and Moco uses OpenSim *Models* to obtain the system’s multibody dynamics, auxiliary dynamics (e.g., muscle activation dynamics and tendon compliance), and kinematic constraints. Moco handles kinematic constraints, which are commonly used to model anatomy such as the knee, shoulder, and neck [48–51].
- **boundary constraints**: Users can enforce average speed, symmetry, or periodicity with constraints relating initial and final states.
- **path constraints**: Users can constrain any function of time to lie in a specified range over the motion. For example, researchers often estimate muscle activity with electromyography and Moco allows constraining simulated muscle excitations to be close to those measurements via the *MocoControlBoundConstraint* class. (These path constraints are not related to OpenSim’s actuator paths.)
- **parameter optimization**: Users can optimize model properties, such as a body’s mass, a muscle’s optimal fiber length, or an exoskeleton’s stiffness.
- **bounds on variables**: Users can bound the values of states, controls, and initial and final time.

The modules of a *MocoProblem* can be combined in diverse ways, as demonstrated by the following examples.

- **dynamically-constrained inverse kinematics**: minimize the error between experimental and model marker positions (marker tracking) and control effort while obeying multibody dynamics. The optimized variables are generalized coordinates, speeds, and forces.
- **electromyography-constrained muscle force estimation**: for a prescribed experimental motion, minimize muscle excitations while obeying multibody dynamics and the difference between muscle excitations and electromyography data. The muscle excitations are the only variables in this problem.
- **torso mass calibration**: for a prescribed experimental motion, minimize model residual forces at the pelvis while obeying multibody dynamics. The joint torques and torso mass are the variables.
- **prediction of muscle coordination adaptations to an exoskeleton**: minimize squared muscle excitations and the error between experimental joint angles and model coordinate values (state tracking) while obeying multibody and muscle activation dynamics. The model coordinates, speeds, muscle excitations, muscle activations, and exoskeleton torques are the variables.

*MocoProblem* describes the optimization problem in Eq (1). We seek the time-dependent states *y*(*t*) and controls *x*(*t*) that minimize a sum of costs *J*_*j*_ with weights *w*_*j*_. The states include generalized coordinates *q*(*t*), generalized speeds *u*(*t*), and auxiliary states *z*(*t*), such as muscle activations. We may also seek time-invariant parameters *p*, the initial time of the motion *t*_0_, or the final time of the motion *t*_*f*_. We place lower (*L*) and upper (*U*) bounds on all variables, as well as bounds on the initial and final values of the states and controls; these bounds allow solving boundary value problems such as standing from a squat.

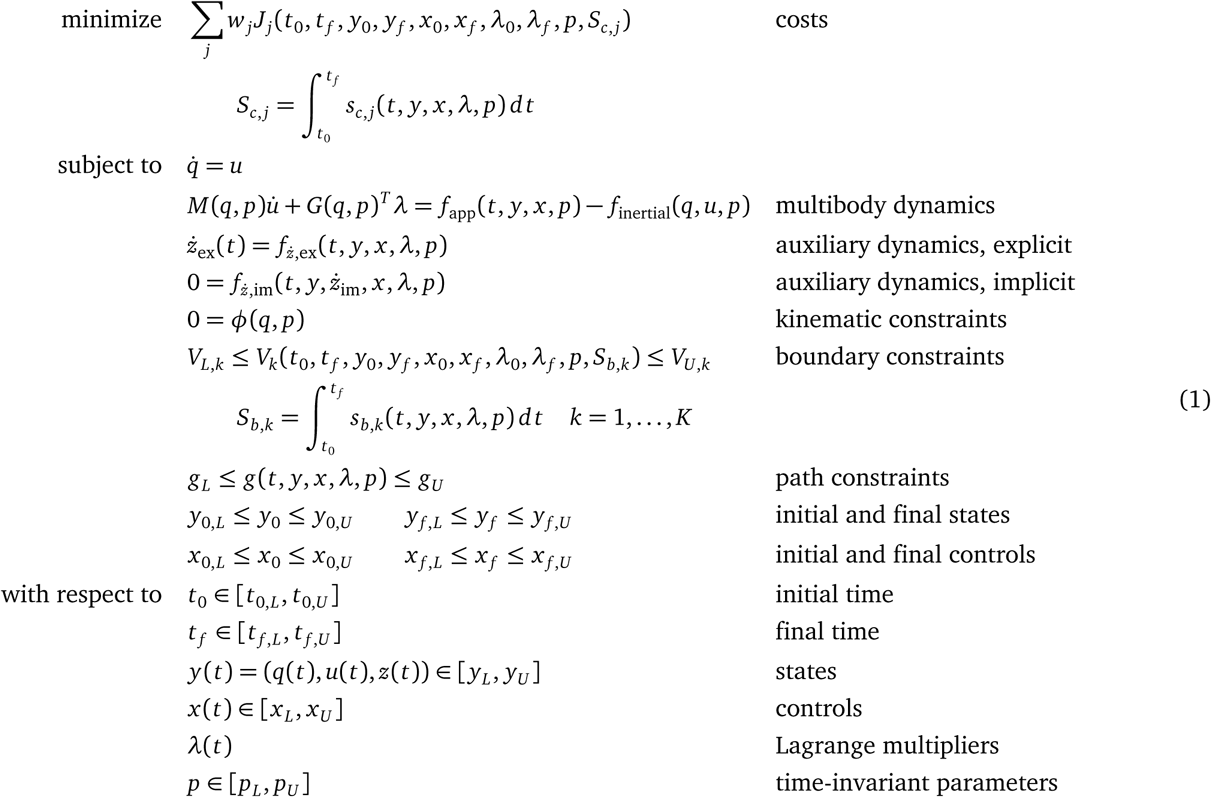

The variables must obey the system’s multibody dynamics (involving the mass matrix *M*; applied forces *f*_app_ from gravity, muscles, etc.; and centripetal and Coriolis terms *f*_inertial_) and any auxiliary dynamics, which may be expressed as explicit 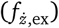 or implicit 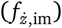 differential equations. The system may contain position-level (holonomic) kinematic constraints to, for example, weld a foot to a bicycle pedal. Each constraint is enforced by forces exerted by tissue, bones, bodies, or other parts of the modeled system. These generalized constraint forces are applied in the constrained directions (e.g., the six degrees of freedom between the foot and pedal), and we introduce time-varying Lagrange multiplier variables *λ* to solve for these forces. The derivative of the kinematic constraints *φ* yields the kinematic constraint Jacobian *G*; the transpose of this matrix converts the Lagrange multipliers into generalized forces along the system’s degrees of freedom. See S1 Appendix for details on how Moco handles kinematic constraints.

Additionally, the variables must obey boundary constraints *V*_*k*_ (with bounds *V*_*L,k*_ and *V*_*U,k*_) and algebraic path constraints *g* over the motion (with time-invariant bounds *g*_*L*_ and *g*_*U*_). The cost terms and boundary constraints may depend on initial and final time; states; controls; kinematic constraint multipliers (required for joint reactions); time-invariant parameters; and an integral, *S*_*c,j*_ or *S*_*b,k*_, over the motion.

We demonstrate Moco’s interface with a MATLAB example that seeks the force to apply to a point mass to move the mass by one meter (starting and ending at rest) in minimum time:

~~~
study = MocoStudy();
problem = study.updProblem();
problem.addGoal(MocoFinalTimeGoal()); % minimum-time problem
problem.setModel(Model(’sliding_mass.osim’)); % load model from file
problem.setTimeBounds(0, [0, 5]); % initial time = 0, final time <= 5 s
problem.setStateInfo(’/slider/position/value’, [-5, 5], 0, 1); % move 1 m
problem.setStateInfo(’/slider/position/speed’, [-50, 50], 0, 0);
problem.setControlInfo(’/actuator’, [-50, 50]);
solution = study.solve();
~~~

Setting the model and adding a cost (termed “goal” in the example) each require only a single statement. Setting bounds on states and controls by name is easier and less bug-prone than setting bounds by index, as is common in other direct collocation software.

Direct collocation relies on gradient-based optimization, and therefore converges faster and more reliably when all functions in the optimal control problem are continuous and differentiable. To this end, Moco includes a muscle model *DeGroote-Fregly2016Muscle* [37] and a compliant contact force model *SmoothSphereHalfSpaceForce* [52] that are continuous and differentiable.

Users wishing to employ a cost term, boundary constraint, or path constraint that Moco does not support can create a C++ plugin using the same steps as for OpenSim plugins. By providing a library of cost, boundary constraint, and path constraint modules, allowing users to create their own modules, and allowing these modules to be combined, we achieve our design goals of ease-of-use, customizability, and extensibility.

### Solving problems with *MocoSolver*

All details of solving an optimal control problem are encapsulated in *MocoSolver*, which is decoupled from *MocoProblem* for flexibility. The *MocoProblem* knows nothing about the solver that may be used. When a user defines a custom cost term, they need not worry about how the solver will handle the cost. The only assumption made by *MocoSolver* about the problem is that it describes a multibody system; this allows using special solver algorithms not suited to generic dynamic systems, such as for handling kinematic constraints [53]. Moco users can add bodies or muscles to their model without modifying the solver, a convenience often not afforded by custom research code that couples the problem formulation to the solver. *MocoSolver* uses the CasADi library [54] to transcribe the continuous optimal control problem defined by *MocoProblem* into a finite-dimensional nonlinear program, which we solve with well-established gradient-based nonlinear program solvers such as IPOPT [55] and SNOPT [56] (see S1 Appendix).

Moco provides two transcription schemes: the second-order trapezoidal scheme and the third-order Hermite-Simpson scheme [17]. Multibody dynamics can be expressed with either explicit differential equations (“forward dynamics”) or implicit differential equations (“inverse dynamics”); problems may converge faster in implicit mode [36, 37]. Solving a musculoskeletal optimal control problem often requires trying many problem formulations and solver settings. Moco users can change the transcription scheme, dynamics mode, and other solver settings with a single line of code.

Solving a *MocoStudy* yields a *MocoSolution* (Figure 2), which is a subclass of *MocoTrajectory* and provides easy access to the values of all variables at any iteration in the optimization. Users provide initial guesses via *MocoTrajectory*, and can use the solution from one problem as the initial guess for a subsequent problem; this permits users to build a complex study by solving a series of simpler studies. For example, a simulation to predict the change in walking kinematics due to an external perturbation (e.g., an ankle exoskeleton) could benefit from an initial guess generated from a series of tracking simulations where kinematic deviations from reference data are gradually penalized less in the objective. *MocoSolution* provides additional information, including whether the solver converged, the final objective value, and the number of solver iterations.

**Figure 2.**
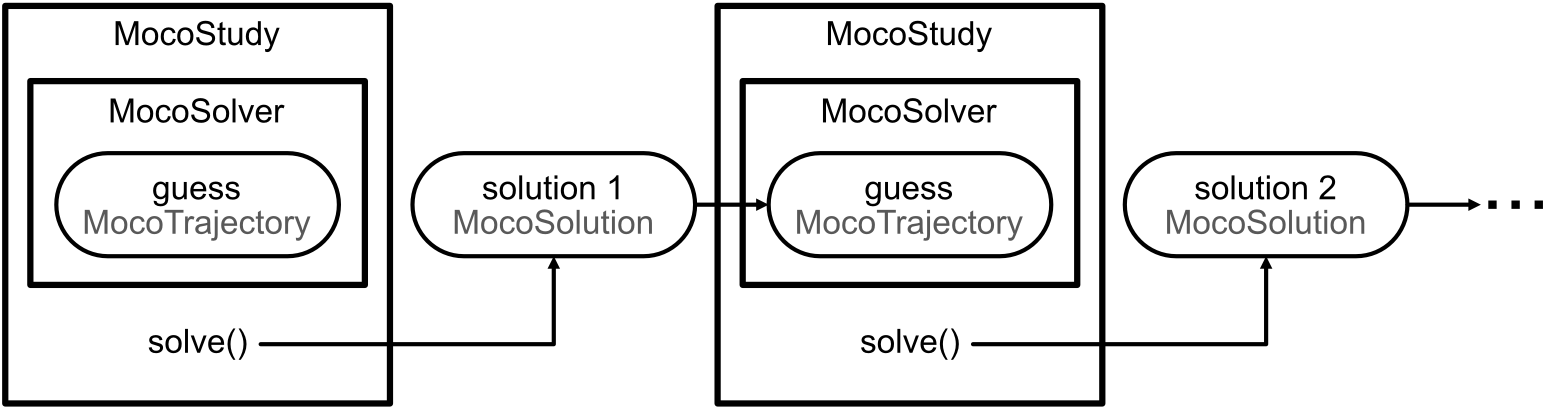
Using trajectories to solve problems iteratively. Guesses for the optimization are specified using *MocoTrajectory*, which holds the values of states, controls, Lagrange multipliers, and parameters at any iteration in the optimization. *MocoSolution* is a subclass of *MocoTrajectory* that holds the solution to a study and includes the success status of the optimization, the final objective value, and the number of solver iterations. Users can use the solution of one problem as the initial guess for a subsequent problem.

After solving a problem, users often wish to visualize the solution as an animation, plot the state and control trajectories, or compute quantities from the solution. With a *MocoStudy* and *MocoSolution*, each of these tasks require only a single line of code. *MocoTrajectories* can be written to and read from tab-delimited *Storage* text files, which are familiar to OpenSim users. The ability to save *MocoTrajectories* and *MocoStudies* to files allows users to reproduce each other’s results, which is essential for sound science [57].

### Tools for standard problems

Currently, Moco provides two tools for solving standard problems (Figure 3):

**Figure 3.**
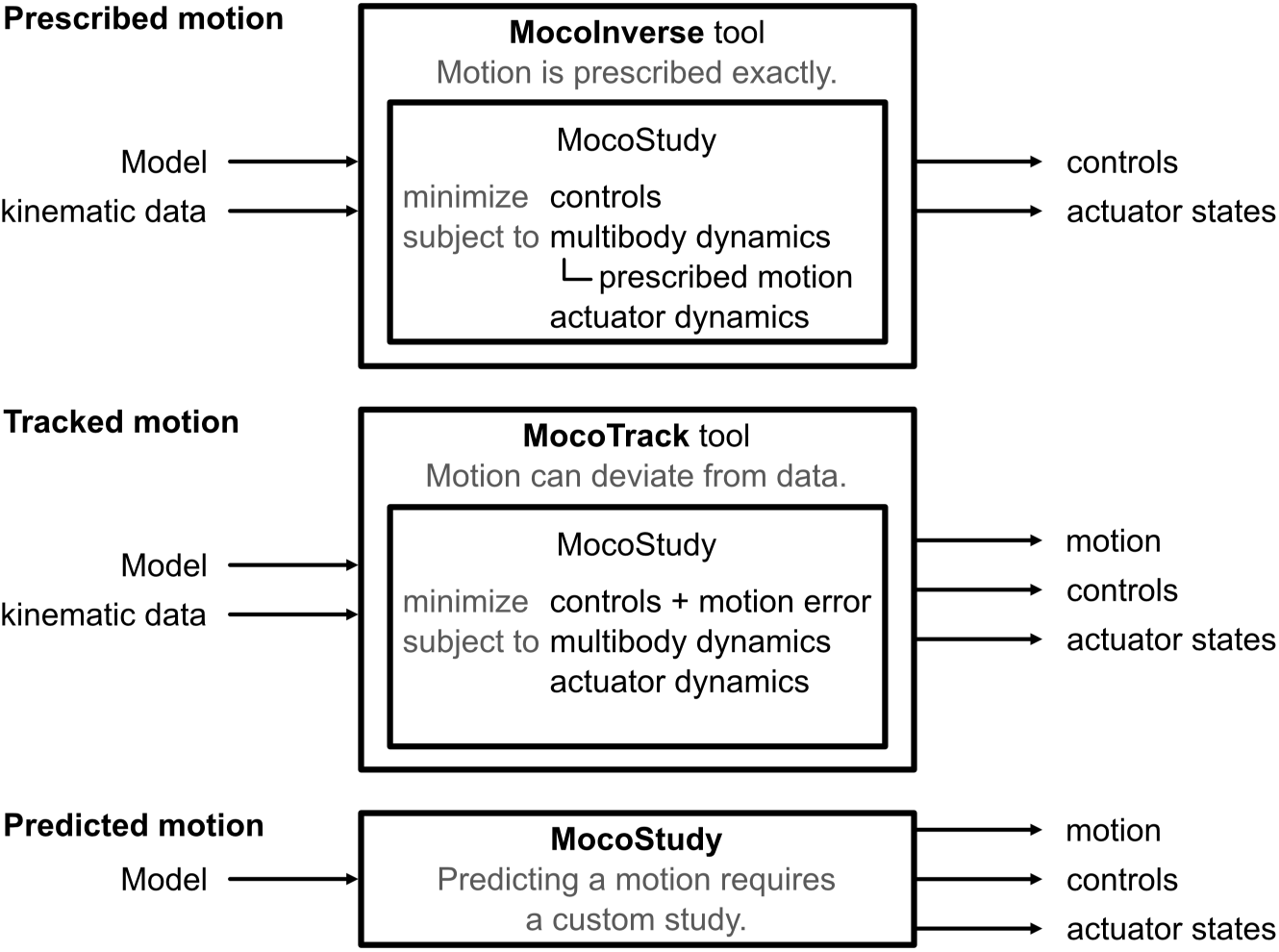
Solving prescribed motion, tracked motion, and predicted motion problems. Moco provides the tools *MocoTrack* and *MocoInverse* for solving standard problems. Both require a *Model* and kinematic data as inputs and produce controls and actuator states as outputs, but these tools solve different optimal control problems. *MocoTrack* produces a new simulated motion, while *MocoInverse* does not permit deviations from the provided kinematic data. Predicting a motion is not easily standardized and requires using a custom *MocoStudy*.

- *MocoInverse* solves the muscle/actuator redundancy problem [37], wherein we solve for muscle (or other actuator) controls that achieve a motion that is prescribed exactly (see S1 Appendix) while minimizing effort or other costs.
- *MocoTrack* solves motion tracking problems, wherein we solve for both a motion and muscle (or other actuator) controls that minimize the error with an observed motion in addition to effort or other costs.

*MocoTrack* is useful for predicting deviations from motion data (e.g., predicting kinematic adaptations to an exoskeleton), while *MocoInverse* is a faster option when the motion should be enforced exactly (e.g., estimating elastic energy storage for an observed motion). *MocoTrack* can use contact models instead of applying measured external forces to the model, as would be done with *MocoInverse*. Applying measured external forces often requires introducing non-physiological “residual” actuators to resolve inconsistencies between measured forces, measured kinematics, and mass properties. For both tools, the only required inputs are an OpenSim model and motion data (coordinate or marker trajectories, and external forces). Internally, the tools build a *MocoStudy* with solver settings that yield fast and reliable convergence on problems we tested.

Future versions of Moco may include tools for model calibration, electromyography-driven simulation, and other standard problems. For problems that do not fit into a standard form, such as predicting a motion, using *MocoStudy* provides the necessary flexibility.

### Verification

Verifying software is essential for good science and gaining the trust of users [58]; thus, we conducted extensive verification tests. For example, producing the known solution to an optimal control problem verifies that Moco implements direct collocation correctly. We solved the following linear optimal control problem, which has a known solution:

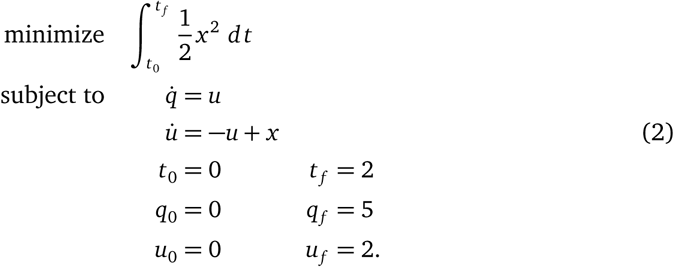

Moco’s solution for the optimal control matched the known control solution with a root-mean-square error of 8.0×10^*−*8^.

Next, we ensured that a time-stepping forward simulation using controls from a motion prediction produced the same motion as in the prediction. These tests used a model consisting of a point mass suspended by three muscles (*DeGrooteFregly2016Muscle*) and under the influence of gravity (Figure 4). For this problem, the muscles had activation dynamics and rigid tendons. We first predicted the state and control trajectories to move the point mass between prescribed initial and final positions, starting and ending at rest (Figure 4, gray band). In this prediction, the cost included both the sum of squared muscle excitations and the final time. Then, we used the predicted controls to perform a time-stepping forward simulation using an OpenSim integrator (Figure 4, blue line). The resulting position trajectory of the point mass matched that from the prediction with a root-mean-square error of 0.0051 m (1.7% of the distance between the initial and final positions). This gives us confidence that Moco enforces the same multibody and muscle dynamics enforced in a time-stepping forward simulation in OpenSim. For more complex problems, conducting a time-stepping forward simulation using controls from a *MocoSolution* requires a stabilizing feedback controller to counteract numerical errors.

**Figure 4.**
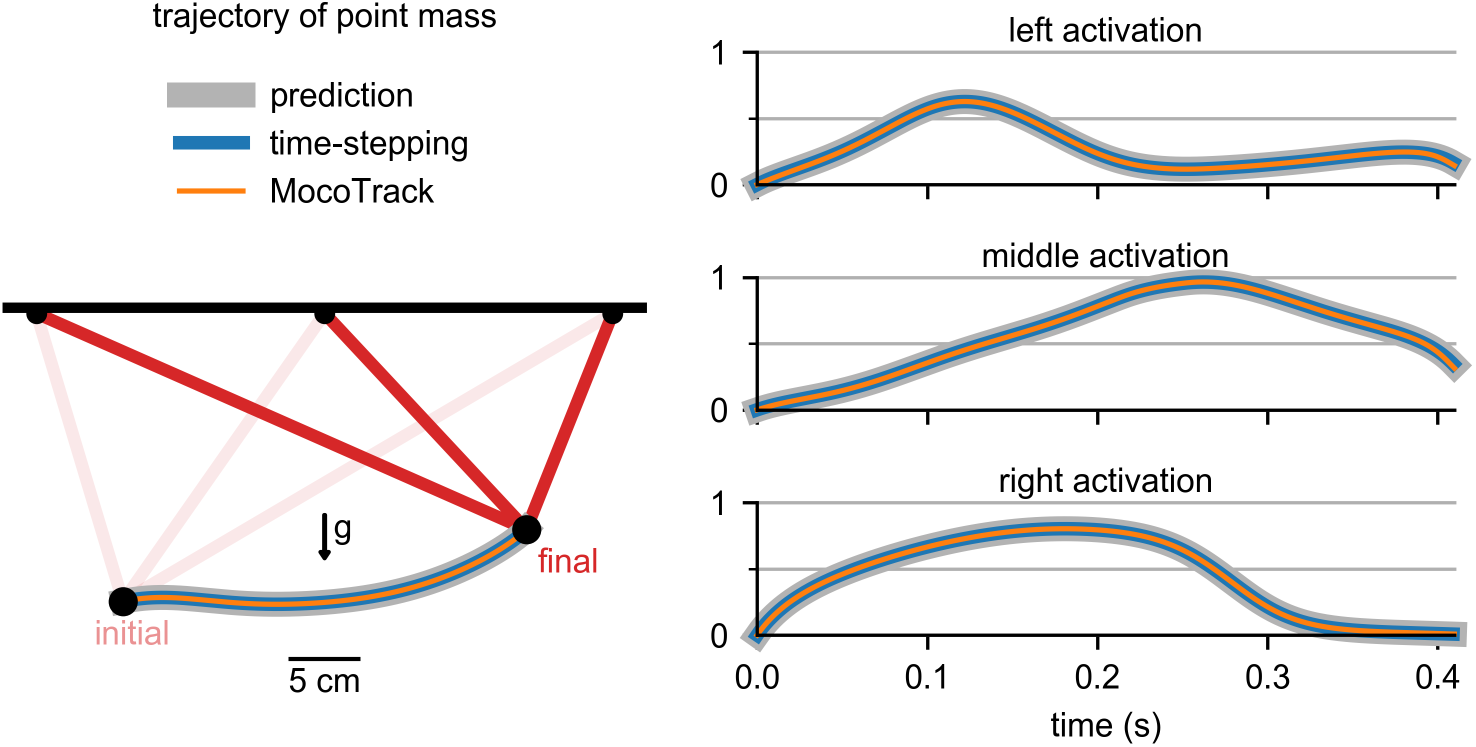
Verification of time-stepping and motion tracking. The trajectory of a point mass suspended by three muscles and moving under the influence of gravity (*g*) in three different types of simulation is shown on the left. The activations of the “left,” “middle,” and “right” muscles throughout the motion are shown on the right. We predicted a trajectory (gray band) that minimized the sum of squared muscle excitations and final time. We then performed a time-stepping forward simulation (blue) with the predicted controls and produced the motion we originally predicted. Tracking the predicted motion with *MocoTrack* (orange) produced the original activations.

To gain confidence in motion tracking problems, we ensured that using *MocoTrack* on a synthesized motion with known muscle activity produced the original muscle activity. We used the same suspended point mass model and tracked the previous motion prediction (Figure 4, gray band) while minimizing squared excitations. The muscle activations from *MocoTrack* (Figure 4, orange line) matched those from the prediction with a root-mean-square error that was 0.23% of the peak predicted activation. To show that there are indeed multiple muscle activity trajectories that produce the same motion, we tracked the motion while minimizing the sum of muscle excitations raised to the fourth power; this resulted in a much larger root-mean-square error that was 11% of the peak predicted activation. For most tracking problems of interest, we do not know the true muscle activity solution; verifying tracking problems using synthesized data gives us confidence in Moco’s estimates of muscle activity for real-world data.

Moco contains an automated test suite with over 30 files that extends beyond the verification described here. For example, the test suite ensures that users receive error messages for incorrect input, that a *MocoStudy* can be written to and read from an XML file, that kinematic constraints are enforced, and that implicit and explicit differential equations lead to the same solution. We ensure all these tests succeed before accepting changes to the code.

## Results

### Estimating muscle activity from motion capture data of walking

Moco can estimate muscle activity in walking, which allows studies of gait disorders and muscle coordination. We used a model with 19 degrees of freedom and 80 lower-limb muscles [50] to simulate one gait cycle of walking at a self-selected speed of 1.25 m/s. The scaled model and data are based on the supplementary material of [50]. All muscles had activation dynamics but only the gastrocnemii and soleus had compliant tendons [37]. We solved for muscle activity using *MocoInverse*, which prescribes kinematics exactly (see “Tools for standard problems” for details), and compared the resulting muscle activations to electromyography measurements (Figure 5).

**Figure 5.**
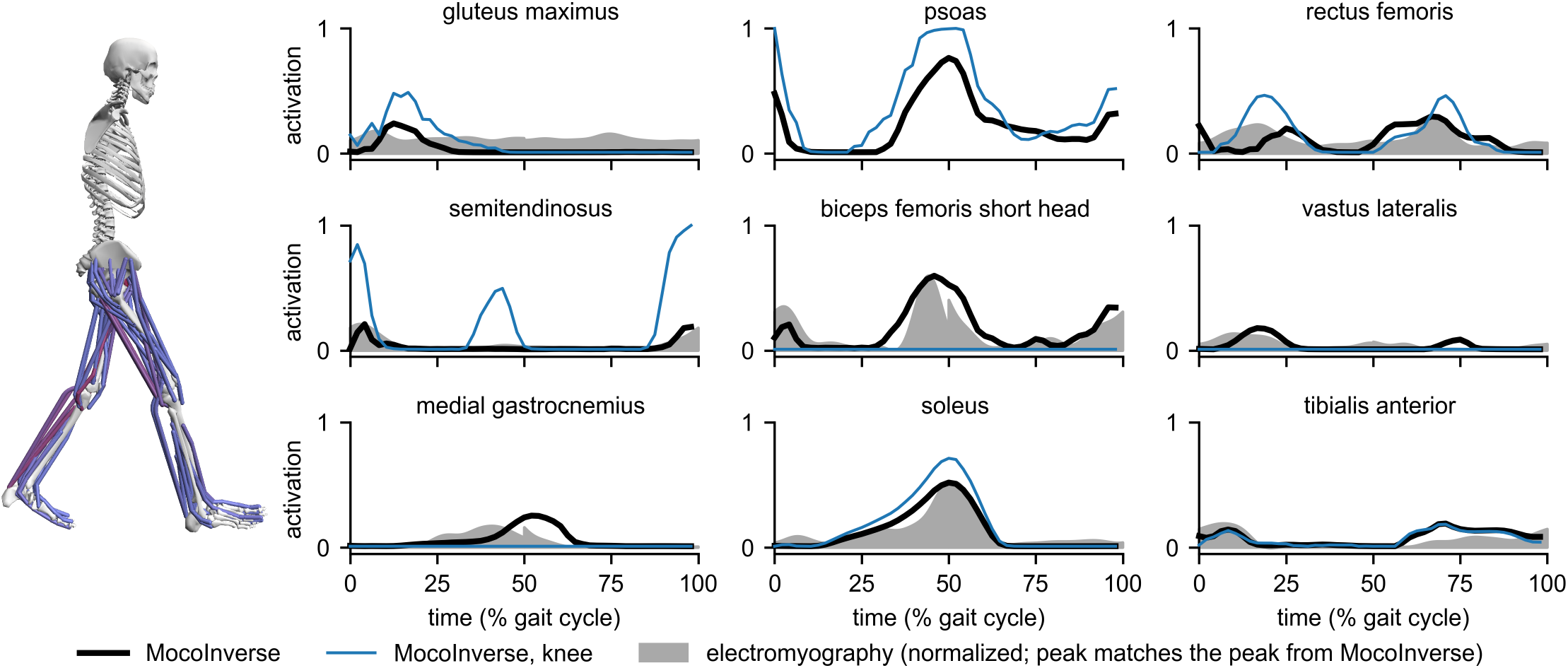
Estimates of muscle activity during walking. *MocoInverse* produced muscle activations (black) whose timing matched the timing from electromyography measurements (gray; available for all muscles shown except psoas) [59]. Electromyography was normalized such that its peak matched the peak of the *MocoInverse* activations. Minimizing knee joint loading reduced the activity of the vastus lateralis, biceps femoris short head, and medial gastrocnemius, which span the knee (blue).

*MocoInverse* produced activations that included some of the major features of the electromyography data [59], such as the timing of peak activity for the semitendinosus, biceps femoris short head, vastus lateralis, medial gastrocnemius, and soleus. With *MocoInverse*, the difference between required net joint moments and muscle-generated net joint moments (termed “reserve” moments in OpenSim) had a maximum value of 2.5 N-m across time and degrees of freedom [58]. *MocoInverse* solved this problem in 4.4 minutes, using a 3.6 GHz Intel Core i7 processor with 8 parallel threads; this duration is similar to the amount of time required by the Computed Muscle Control [9] tool in OpenSim for solving prescribed motion problems [50].

To demonstrate how Moco allows customization, we added a cost term to *MocoInverse* to minimize knee joint loading. With this additional cost, the peak magnitude of the knee joint reaction force decreased from 4.7 to 2.9 body weights. As expected, the activity of muscles crossing the knee joint (vastus lateralis, biceps femoris short head, and medial gastrocnemius) decreased. To compensate for the reduced medial gastrocnemius moment at the ankle, soleus activity increased, as seen in a previous simulation study [60]. Activity of the semitendinosus increased to compensate for the decrease in activity of the biceps femoris short head. Psoas activity increased to its maximum value; while a subject attempting to minimize the same objective in an experiment may not maximally activate their psoas, this high activity was allowed by our problem definition. Joint loading is a more complex cost than simply minimizing excitations; the problem solved in 48 minutes.

### Predicting and assisting a squat-to-stand motion

To show that Moco can predict motions and optimize parameters of a model, we predicted a squat-to-stand motion that minimized a combination of effort, expressed as the sum of squared excitations, and the duration of the motion. The initial pose was prescribed to be squatting [4], and the final pose was prescribed to be upright standing. No motion was tracked. The model contained a torso and leg with 9 muscles obeying activation dynamics and compliant tendon dynamics. To enforce mediolateral symmetry, we used a single leg and doubled muscle strengths, which we obtained from a previous study [14]. We used Moco’s default initial guess, in which each variable’s value is the midpoint of the variable’s bounds. The predicted motion and muscle activations are shown in Figure 6. As expected, extensor muscles such as the gluteus maximus, hamstrings, and vasti exhibited the greatest activity.

**Figure 6.**
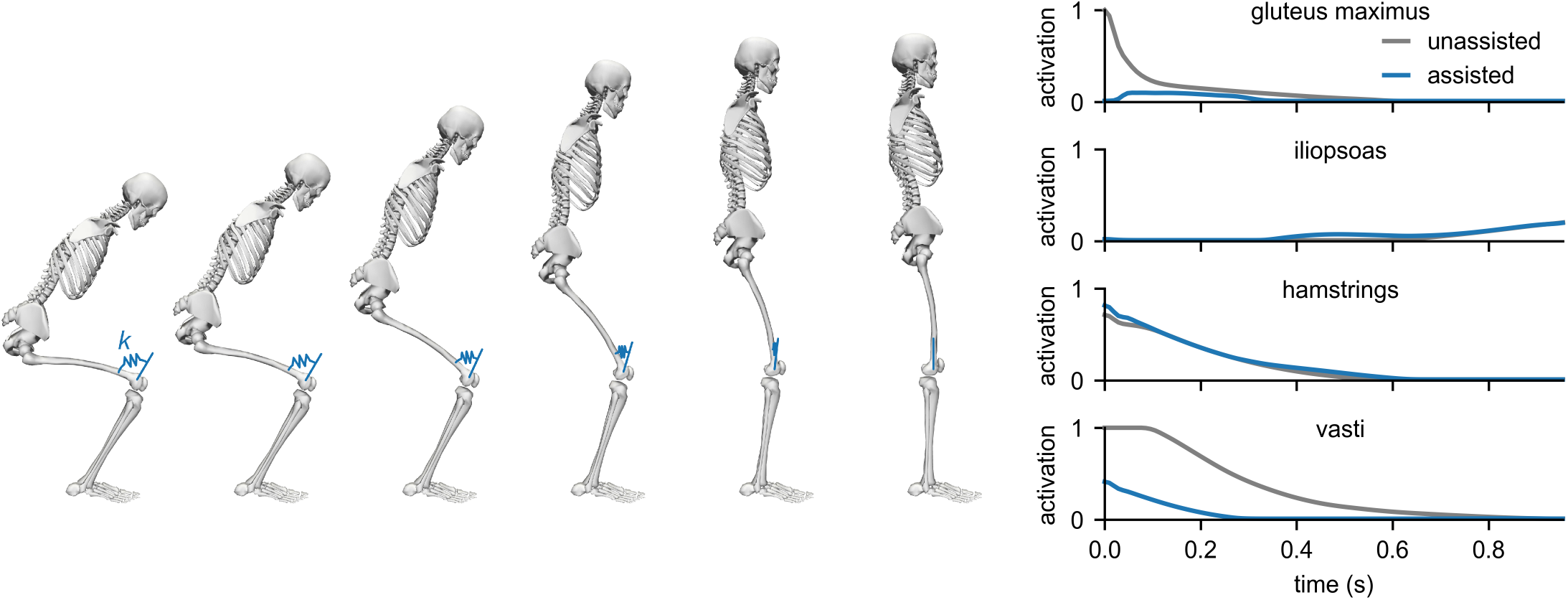
Predicting and assisting a squat-to-stand motion. We predicted a squat-to-stand motion with prescribed initial and final poses that minimized the sum of squared muscle excitations and final time (gray). The predicted motion is shown on the left, and the activations of key muscles are shown on the right. We then added a torsional spring to the knee and solved for the optimal motion, muscle activations, and spring stiffness *k* (blue). The spring allowed gluteus maximus and vasti activity to decrease substantially.

Next, we added a torsional spring to the knee and solved for the optimal motion, muscle activations, and spring stiffness. The spring was in equilibrium when the knee was extended, as illustrated in Figure 6. We used the same cost as in the unassisted case; no motion was tracked. With the assistive device, the motion was achieved with lower muscle activity. The optimal spring stiffness was 88 N-m/rad.

Both predictions solved in under 3 minutes. Moco’s ability to rapidly predict motions and optimize device parameters makes it a valuable tool for designing assistive devices.

### Availability and future directions

OpenSim Moco can be downloaded freely for Windows and Mac from SimTK and GitHub (https://simtk.org/projects/opensim-moco and https://github.com/opensim-org/opensim-moco), where we develop the project and users can report bugs and request features. The OpenSim Moco source code is available under the permissive Apache License 2.0, though some dependencies have more restrictive licenses (e.g., CasADi [54] is available under the GNU Lesser General Public License).

The documentation for Moco contains a User Guide, Theory Guide, Developer Guide, and an Application Programming Interface (API) Reference. The User Guide explains how to use Moco and provides tips for posing a problem. The Theory Guide explains how Moco implements direct collocation, and the Developer Guide introduces the code and explains software design choices. The API Reference describes the classes and functions in the library. Lastly, we provide a printable two-page “cheat sheet” that demonstrates common commands in Moco.

The Moco distribution contains examples in MATLAB, Python, and C++. These examples range from predicting the optimal trajectory for a double pendulum to predicting 2-D muscle-driven walking (which solves in under an hour). The code used to generate the results for this paper are available at https://github.com/stanfordnmbl/mocopaper.

Moco currently lacks the ability to handle certain problems. Metabolic energy consumption is a commonly used cost term, but direct collocation struggles with the non-convexity of many energy consumption models. Koelewijn et al. recently published a smoothed energy consumption model for use in direct collocation [39]; we hope Moco will include this model in the future. Implementing a cost term that minimizes the error between measured and simulated contact forces would improve the accuracy of simulated ground reaction forces in tracking problems. Many direct collocation formulations allow a problem to contain multiple phases, each with different system dynamics. Supporting multiple phases enables modeling foot–ground contact with kinematic constraints instead of compliant contact, which would avoid the poor numerical conditioning caused by the stiffness of compliant contact. Falisse et al. used a multiple-phase approach to solve for muscle parameters that fit multiple unrelated motions [34]. By including cost terms for energy expenditure and contact force tracking and permitting multiple phases, Moco could cover a wider range of biomechanics applications.

The performance and ease of use of Moco’s direct collocation solvers could be improved. Supporting mesh refinement would allow the solver to increase the number of mesh intervals in time ranges with fast dynamics, thereby improving accuracy. Computing the nonlinear program derivatives with algorithmic differentiation instead of finite differences would vastly improve the speed of Moco, but would require substantial changes to the source code of OpenSim and its dependencies [40].

We designed Moco to be easy to use, customizable, and extensible. We verified the software and applied it to multiple musculoskeletal problems. Given this foundation, we expect Moco to accelerate research by reducing the time spent wrestling with simulation tools and enabling our field to tackle more ambitious problems. Moco handles models with kinematic constraints, muscle activation dynamics, compliant tendons, and compliant contact, and can minimize a combination of complex costs such as marker tracking and joint reaction loads. To build a community that sustains the Moco project, we have organized several in-person workshops. We invite the community to help improve the software and contribute documentation, examples, teaching materials, and code.

## S1 Appendix

### Kinematic constraints

Support for kinematic constraints is a key feature of Moco. Understanding how kinematic constraints are handled in multibody dynamics is valuable for understanding how Moco handles such problems. Consider a two-dimensional point mass system with coordinates *q*_*x*_ and *q*_*y*_ constrained to a parabola 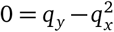. To prevent the point mass from violating the constraint, we must apply a force perpendicular to the parabola. Each constraint has a corresponding scalar force variable, called a Lagrange multiplier *λ*. We must solve for the required magnitudes of these constraint forces, but the direction in which we apply each of these forces is determined by the derivative of the constraint equations. We gather the derivatives of the constraint equations in the kinematic constraint Jacobian matrix *G*. Each row in this matrix contains the derivative of a single constraint equation with respect to each degree of freedom, and the matrix has a column for each degree of freedom. For the parabola example, the Jacobian is (−2*q*_*x*_,1). The transpose of this matrix, *G^T^*, contains columns that are vectors in the state space which are perpendicular to each constraint. For our single constraint, the vector (−2*q*_*x*_,1)^*T*^ is perpendicular to the parabola. The matrix-vector product between the Jacobian transpose and the Lagrange multipliers, *G*^*T*^ *λ*, yields the vector of generalized forces (whose length is the number of degrees of freedom) necessary for enforcing the system constraints. For our point mass example, the generalized forces yielded by the vector-scalar product (−2*q*_*x*_,1)^*T*^ *λ* keep the point mass on the parabola. To apply these forces to the multibody system, we include the *G*^*T*^ *λ* term in the multibody dynamics equations of motion.

When simulating a multibody system with time-stepping forward integration, we first ensure the initial generalized coordinates and speeds satisfy the kinematic constraints *ϕ*(*q*) = 0 and their first derivative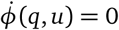 via a root-solve. During the integration, we solve for generalized accelerations and Lagrange multipliers that obey the multibody dynamics equations of motion and the second derivative of the kinematic constraints, 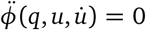. Numerically integrating the resulting generalized accelerations yields generalized coordinates and speeds that approximately lie on the constraint manifold defined by *ϕ*(*q*) = 0; to fix any errors in the constraints caused by numerical integration error, we project the generalized coordinates and speeds back onto the constraint manifold [47].

In direct collocation, the generalized coordinates, generalized speeds, and Lagrange multipliers are the unknowns—we are solving for the entire trajectory of the system. When expressing multibody dynamics as implicit differential equations, the generalized accelerations are also unknowns. How we solve for the trajectory of the system in the presence of kinematic constraints depends on the transcription scheme; see the remainder of this Appendix for details.

### Defining problems containing prescribed kinematics

A common task in musculoskeletal biomechanics is to estimate the muscle and actuator behavior that drove an observed motion. We can solve this problem by minimizing the error between the observed motion and the simulated motion, as with Computed Muscle Control (using the “slow target”) [9] or *MocoTrack*. Alternatively, we can prescribe the motion exactly, as with Static Optimization, electromyography-driven simulation [10], and the Muscle Redundancy Solver [37]. Consider a two-dimensional point mass with coordinates *q*_*x*_ and *q*_*y*_ for which we prescribe a circular motion via the functions 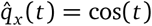 and 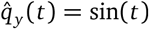. We can either add these functions to the kinematic constraints *ϕ*(*q*), or we can substitute these functions into the equations of motion, thereby eliminating the variables *q*_*x*_ and *q*_*y*_. With Moco, users can choose either the former approach through OpenSim’s *Coordinate*, or the latter (and usually preferable) approach using *PositionMotion*, a new component that employs Simbody’s *Motion* class. Prescribing kinematics by eliminating variables leads to a problem that is robust and fast— the nonlinear multibody dynamics are removed from the optimization problem—but prevents predicting kinematic deviations from the observed motion.

When we prescribe kinematics in Moco by eliminating variables, we replace the problem in Eq (1) with the following:

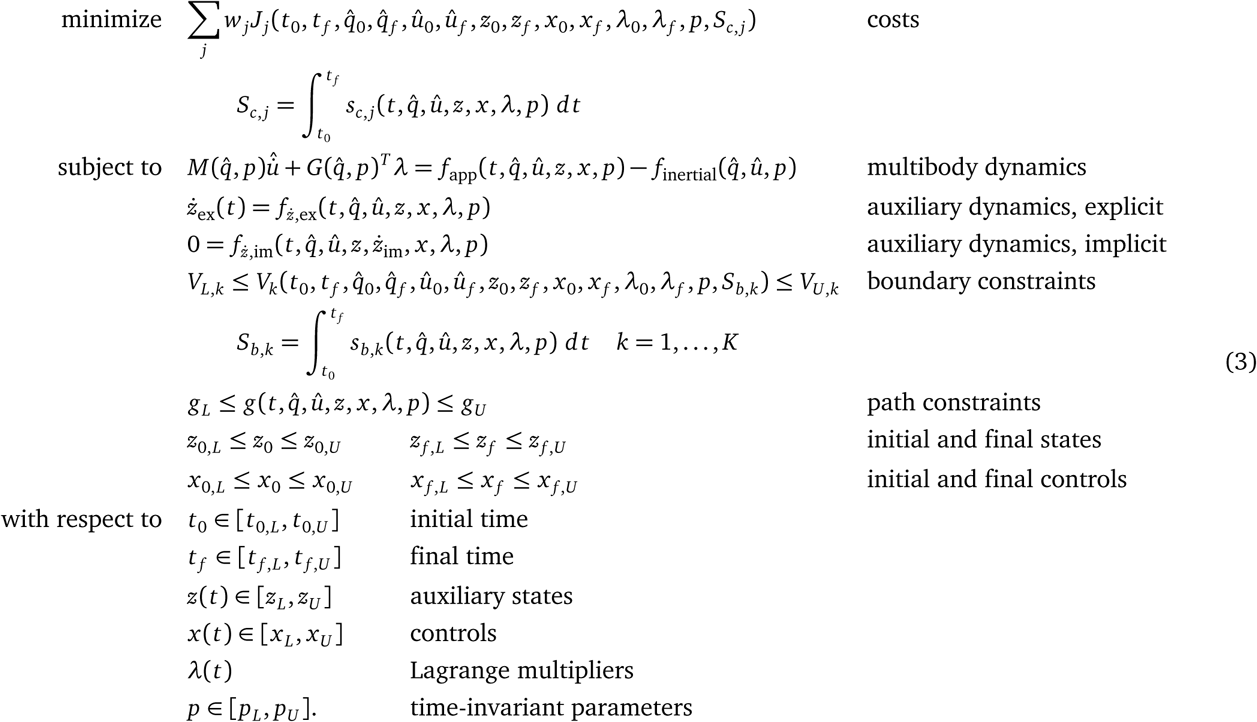

We replace the kinematic variables *q* and *u* with known quantities 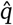 and 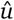. The system still depends on auxiliary state variables *z* and control variables *x*, and includes auxiliary dynamics. If none of the parameter variables affect the multibody system, then the multibody dynamics are reduced to a force balance: muscles and other force elements must generate the net generalized forces determined by the kinematics and external loads data. The easiest way to prescribe kinematics in Moco is to use the *MocoInverse* tool.

Whether the motion is prescribed by adding constraints or eliminating variables, OpenSim supplements the modeled force elements with Lagrange multipliers to ensure the prescribed motion is achieved. When using *PositionMotion* with Moco, we require that the prescribed motion’s Lagrange multipliers are zero, thereby ensuring the motion is fully generated by the modeled force elements.

### Transcription schemes

#### Trapezoidal transcription

The trapezoidal scheme transcribes the optimal control problem into a nonlinear program by approximating integrals using the trapezoidal rule. As a second-order scheme, trapezoidal transcription exhibits accuracy that improves four-fold when halving the mesh interval (i.e., time step).

We discretize the continuous variables *t*, *y*, *x*, and *λ* on a mesh of time points *t*_*i*_ defined by dimensionless time *τ*_*i*_, yielding *n* mesh intervals with durations *h*_*i*_:

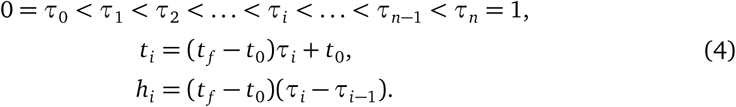

For conciseness, we define the following function:

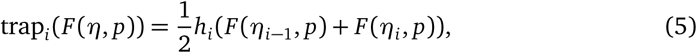

where trap_*i*_() is a trapezoidal rule approximation of the area under the function *F* for mesh interval *i*, and *η* represents any subset of continuous variables. We define the explicit multibody dynamics function as:

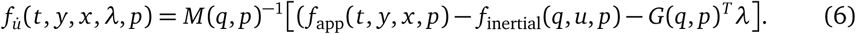

The mass matrix *M*, the centripetal and Coriolis forces *f*_inertial_, and kinematic constraint Jacobian *G* are computed by Simbody (OpenSim’s multibody dynamics engine) using order-N recursive algorithms; the complete mass matrix is not computed explicitly. The applied forces *f*_app_ are usually defined by OpenSim components that implement Simbody force elements. Simbody computes the constraint Jacobian *G* based on any Simbody kinematic constraints that the OpenSim model adds to the system. In Moco, the Lagrange multipliers *λ* are explicit optimization variables; this approach differs from that in forward simulations, in which Simbody solves for accelerations and multipliers simultaneously.

The result of the trapezoidal transcription, with multibody dynamics expressed as explicit differential equations, is the following nonlinear program:

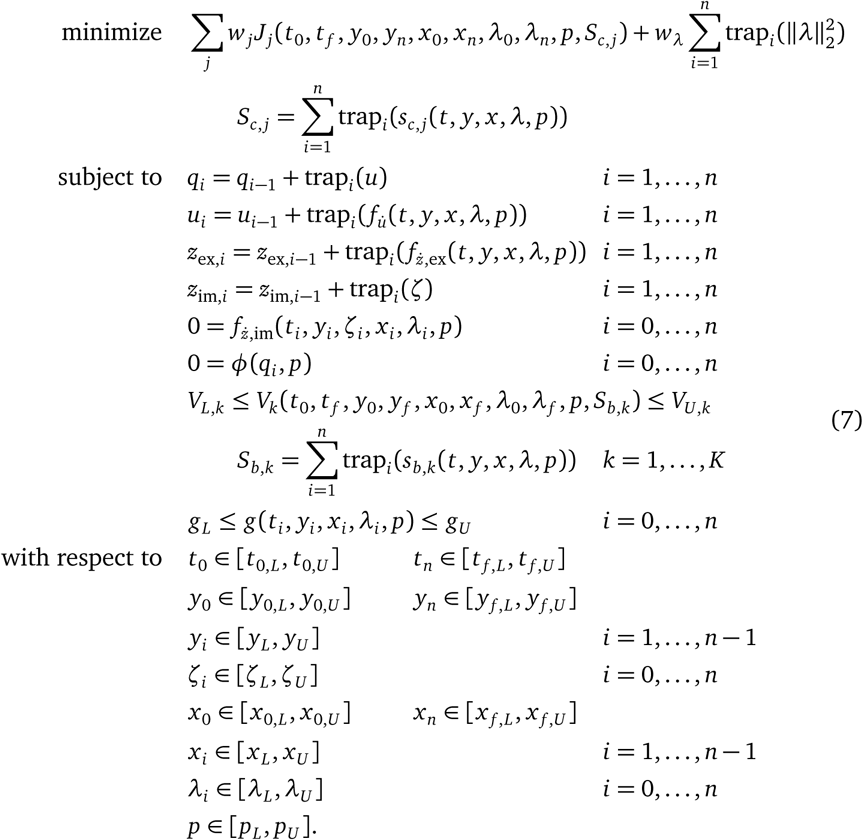

In this form, the problem can be solved directly by a nonlinear program solver. We introduce the algebraic (control) variable *ζ* as the derivative of auxiliary state variables whose dynamics are expressed with an implicit differential equation.

When expressing the multibody dynamics implicitly, we remove the constraint involving 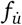, introduce generalized accelerations as an algebraic variable *υ*, and enforce multibody dynamics in “inverse dynamics” form:

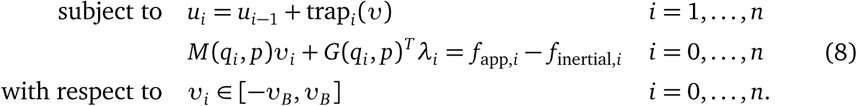

The constant *υ*_*B*_ is a large positive number (1000 by default).

The dynamic, kinematic, and path constraints are enforced at a set of discrete time points, so the quadratic spline approximation to the continuous variables may violate the original continuous-time constraints between the discrete time points. For this reason, a mesh with more points leads to a more accurate solution.

Our implementation of trapezoidal transcription handles kinematic constraints, but not in the most robust fashion. We enforce *ϕ* but not its time derivatives; enforcing the constraints at only the position level yields an index-3 differential-algebraic system of equations, which is challenging to solve [17, 22, 61]. Furthermore, we minimize these multipliers (with weight *w*_*λ*_) to improve numerical conditioning.

When kinematics are prescribed (see “Prescribed kinematics”), multibody dynamics must be expressed implicitly and kinematic constraints are not enforced; we expect the prescribed kinematics to already obey the constraints.

#### Hermite-Simpson transcription

The Hermite-Simpson scheme transcribes the optimal control problem into a nonlinear program by approximating integrals using a Hermite interpolant and Simpson integration rule. As a third-order scheme, Hermite-Simpson transcription exhibits accuracy that improves eight-fold when halving the mesh interval.

We use a similar dimensionless time mesh as for the trapezoidal scheme, with *n* mesh intervals with durations *h*_*i*_. We also introduce collocation points at the midpoints of the mesh intervals, leading to a total of 2*n* + 1 time points at which we discretize the continuous variables:

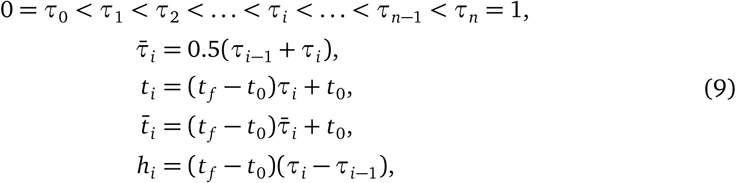

where 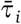 denote mesh interval midpoints. For conciseness, we define the following functions:

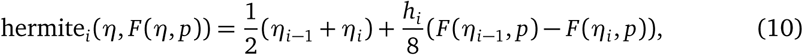

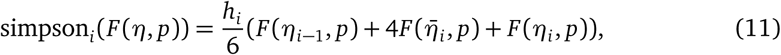

where hermite_*i*_ () represents the Hermite interpolant and simpson_*i*_() represents the Simpson integration rule. Again, *F* is a function for mesh interval *i*, and *η* represents any subset of continuous variables.

Using the explicit multibody dynamics function 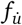 defined previously, Hermite-Simpson transcription results in the following nonlinear program:

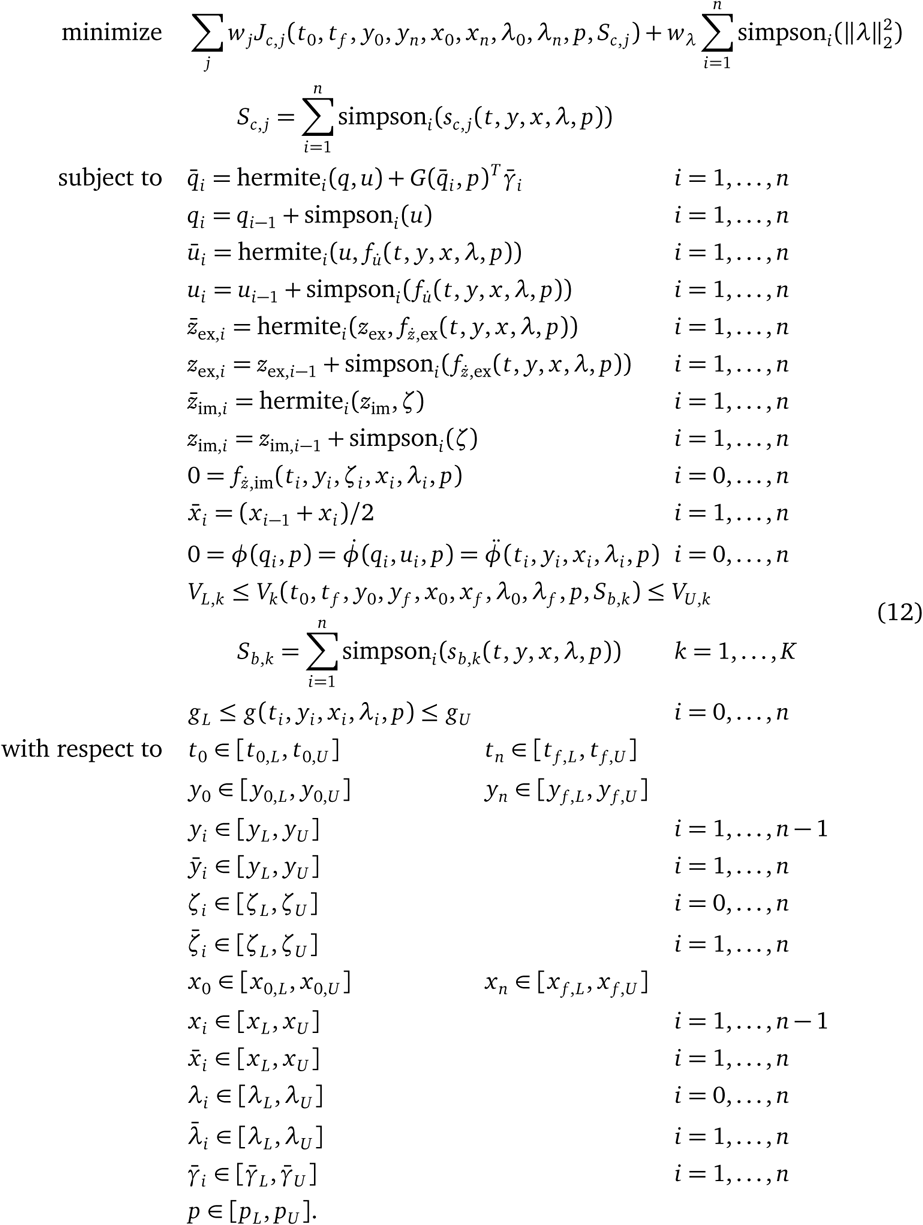

The 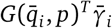 term is a velocity correction that is necessary when enforcing derivatives of kinematic constraints in the nonlinear program. The additional variables 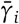 prevent the kinematics from being overconstrained, and the kinematic constraint Jacobian transpose operator 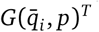 restricts the velocity correction to the tangent plane of the constraint manifold [53]. Currently, we only support enforcing derivatives of position-level, or holonomic, constraints, represented by the equations:

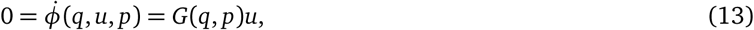

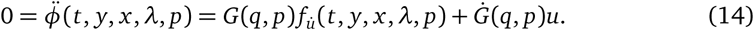

The explicit multibody dynamics function is used here where 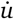 would appear if it were a continuous variable in the problem (as in implicit mode, see below).

Algebraic constraints are not enforced at the midpoints of the mesh intervals, but exhibit fourth-order accuracy at these points [53].

For implicit multibody dynamics, we again remove the constraints involving 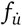 and introduce generalized accelerations as algebraic variables *υ* to enforce multibody dynamics in “inverse dynamics” form:

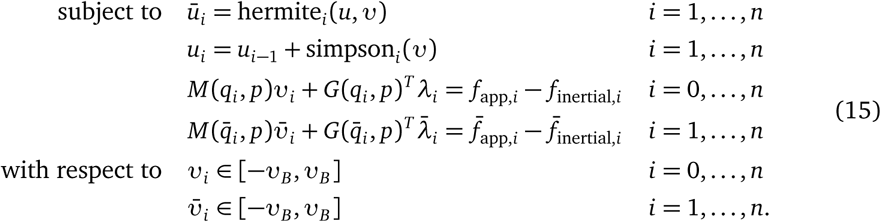

#### Moco’s direct collocation solvers

Moco provides two solvers as subclasses of *MocoSolver*: *MocoCasADiSolver* uses the third-party CasADi library [54], and *MocoTropterSolver* uses a direct collocation solver we developed named Tropter. CasADi is an open-source package for algorithmic differentiation and is a bridge to nonlinear program solvers IPOPT [55], SNOPT [56], and others.

Gradient-based nonlinear program solvers require the gradient of the objective, the Jacobian of the constraints, and sometimes the Hessian of the Lagrangian [17]. To maximize computational efficiency, these derivatives are ideally computed exactly through either analytic expressions or algorithmic differentiation [54, 62]. OpenSim’s main distribution does not provide exact derivatives, so we use finite differences. CasADi is an ideal library for employing direct collocation, but two limitations led us to create Tropter: CasADi did not initially support finite differences, and CasADi’s open-source license is more restrictive than OpenSim’s. More recent versions of CasADi support finite differences and CasADi understands the structure of the nonlinear program objective and constraint functions, allowing for potentially more efficient finite difference calculations than with Tropter, which treats the nonlinear program objective and constraints as black-box functions [63]. If OpenSim provides exact derivatives in the future, we can exploit the algorithmic differentiation modes in Tropter and CasADi [40]. Those distributing Moco as a dependency of closed-source software may prefer distributing Moco without CasADi, as CasADi’s “weak copyleft” GNU Lesser General Public License 3.0 places requirements on how CasADi is redistributed.

## Acknowledgments

We thank Ajay Seth, Michael Sherman, Friedl de Groote, Michael Posa, Joris Gillis, and Joel Andersson for discussing methodology and implementation; Bradley Humphreys, Carmichael Ong, Noah Gordon, and Jenny Yong for contributing code; Andrew Baines, Mohammad Shourijeh, and Prasanna Sritharan for testing the software; and Ayman Habib for reviewing the manuscript. This work was supported by the National Institutes of Health grants U54 EB020405 and P2C HD065690. CLD and NAB received support from the National Science Foundation Graduate Research Fellowship Program. CLD received support from the Stanford Bio-X Graduate Fellowship. NAB received support from the Stanford Graduate Fellowship Program.

